# Test–Retest Reliability of Meta Analytic Networks During Naturalistic Viewing

**DOI:** 10.1101/2024.05.15.594266

**Authors:** Jean-Philippe Kröll, Patrick Friedrich, Xuan Li, Yulia Nurislamova, Nevena Kraljevic, Anna Geiger, Julia Mans, Laura Waite, Julian Caspers, Xing Qian, Michael WL Chee, Juan Helen Zhou, Simon Eickhoff, Susanne Weis

## Abstract

Functional connectivity analyses have given considerable insights into human brain function and organization. As research moves towards clinical application, test-retest reliability has become a main focus of the field. So far, the majority of studies have relied on resting-state paradigms to examine brain connectivity, based on its low demand and ease of implementation. However, the reliability of resting-state measures is mostly moderate, potentially due to its unconstrained nature. Recently, naturalistic viewing paradigms have gained popularity because they probe the human brain under more ecologically valid conditions, thereby possibly increasing reliability. Therefore, we here compared the reliability of graph metrics extracted from resting-state and naturalistic viewing in functional networks, across two sessions. We show that naturalistic viewing can increase reliability over resting-state, but that its effect varies between stimuli and networks. Furthermore, we demonstrate that the effect of naturalistic viewing differs between two cohorts with Asian and European cultural backgrounds. Taken together, our study encourages the use of naturalistic viewing to increase reliability, but emphasizes the need to carefully select the appropriate stimulus and network for the respective research question.

## Introduction

Functional magnetic resonance imaging (fMRI) data has become a widely-used tool to investigate neurological diseases and their underlying patterns (Balthazar et al., 2014; Basaia et al., 2019; Supekar et al., 2008; Wu et al., 2009). The critical assumption behind all of these studies is that the measured brain activity is reliable, such that differences between subjects and timepoints are interpretable. However, the reported reliability of fMRI measures varies vastly across studies (Bennett and Miller, 2010), due to small test-retest samples and different analysis choices. As fMRI research moves towards the identification of biomarkers (Bassett et al., 2008; Rubinov et al., 2009; Supekar et al., 2008; Wang et al., 2009), increasing reliability has become a priority. In order to aid in the diagnosis and prognosis of brain disorders, a measure has to be capable of giving consistent results, otherwise it is unsuitable as a biomarker.

The majority of prior reliability studies have relied on metrics derived from resting state (RS) (Braun et al., 2012; Guo et al., 2012; Wang et al., 2011). With low demands on the participants, RS is well suited for healthy as well as patient cohorts and allows for a quick data acquisition. Although various studies reported moderate to good reliability of RS-derived measures (Braun et al., 2012; Deuker et al., 2009; Wang et al., 2011), the RS paradigm also suffers from a few drawbacks. Data acquired during the RS can be strongly confounded by head movement and drowsiness of the participant due to its unconstrained nature (Tagliazucchi and Laufs, 2014; Van Dijk et al., 2012), as participants struggle to remain awake and motionless in the absence of a task or stimulus. For the same reasons, RS is more susceptible to be influenced by spontaneous thought of the participant (Christoff et al., 2004; Gonzalez-Castillo et al., 2021).

Naturalistic Viewing (NV) paradigms, during which participants are presented with a story or a film, have recently gained popularity because they might give insight into the brain’s function under more ecologically valid conditions. It has been shown that NV poses several advantages over conventional RS such as increased participant engagement, reduced head movement and increased synchronization between subjects (Hasson et al., 2004; Wang et al., 2017). Especially relevant for clinical studies, NV shares with RS the advantage of minimizing demand on the participants (Eickhoff et al., 2020). On the other hand, NV paradigms place a behavioral constraint that allows for the study of normal and abnormal brain function, somewhat similar to task-based designs. Making use of these advantages, a series of studies could show altered connectivity during NV in patients (Guo et al., 2016, 2015; Hyett et al., 2015; Yang et al., 2020), encouraging the application of NV measures as biomarkers.

Furthermore, several studies suggest that NV increases test-retest reliability in comparison with RS (O’Connor et al., 2017; Wang et al., 2017; Zhang et al., 2022). This improvement can be attributed to several factors. First, many studies have pointed out that NV improves signal properties by increasing participant engagement (Eickhoff et al., 2020; Finn and Bandettini, 2020; Li et al., 2022; Vanderwal et al., 2017). Secondly, by reducing head movement and drowsiness, NV is less susceptible to noise than conventional RS. Thirdly, by presenting the same stimulus across sessions, NV is less influenced by spontaneous thought of the participant while also placing a behavioral constraint that reduces variance. However, the effect of NV on reliability is dependent on various factors such as attention (Ki et al., 2016), successful episodic encoding (Hasson et al., 2008) as well as the chosen movie stimulus (Hasson et al., 2010; Kröll et al., 2023; Tian et al., 2021) and differs between different brain regions and networks (Wang et al., 2017).

The present study aims to further evaluate the test-retest reliability of NV, by investigating its influence on the reliability of five commonly used graph theoretical measures. The application of graph theoretical measures to fMRI data is an established method (Braun et al., 2012; Guo et al., 2012; Reijneveld et al., 2007; Stam and Reijneveld, 2007), and has given insights into the complex functional structure of the brain (Bullmore and Sporns, 2009; Rubinov and Sporns, 2010), both in healthy and patient cohorts. To benchmark the reliability of NV, we compare it to that of RS. Further, we evaluate the influence of the movie content, by employing stimuli with different levels of social content, ranging from the neutral movie *Inscapes*, over the silent movie *The Circus*, to the most social movie *Indiana Jones and the Temple of Doom*. In addition, several authors have suggested that the same NV stimuli might deviate in its effect between different populations (Eickhoff et al., 2020; Hasson et al., 2010; Telesford et al., 2010). The cultural background of a participant is likely to influence how a given movie is perceived and might result in deviating effects across cohorts. Therefore, in this study, we compare the effect of NV in two independent samples from Europe and Asia, respectively, using the same stimuli. In contrast to the majority of previous studies, we here compare reliability on the basis of a priori defined networks, and not on a whole-brain basis.

The analysis of network based measures allows us to investigate how NV influences the reliability in different cognitive domains. The networks implemented in this study are meta-analytically defined networks that represent the most likely core nodes involved in a given cognitive function, because they incorporate convergent information from a multitude of studies.

## 2 Methods

### 2.1 Participants

#### Dataset IMAX

For the first dataset, 36 healthy right-handed and ambidextrous adults were scanned at the Centre for Translational MR Research, National University of Singapore. Exclusion criteria were neurological or psychiatric diagnoses, significant visual or hearing impairment, alcohol or caffeine consumption 6 hours prior to the scan and self-reporting of bad sleep the night before the scan days. All participants underwent three identical testing sessions within a one-week interval. Subjects gave written, informed consent and were compensated for their participation. The study was approved by the institutional review board of the National University of Singapore.

#### Dataset JUMAX

For the second dataset, 36 healthy adults were scanned at the Forschungszentrum Jülich. Exclusion criteria were neurological or psychiatric diagnoses, significant visual or hearing impairment, alcohol or caffeine consumption 6 hours prior to the scan and self-reporting of bad sleep the night before the scan days. All participants underwent three identical testing sessions within a one-week interval. Subjects gave written, informed consent and were compensated for their participation. The study was approved by the ethics committee of the Heinrich Heine University, Düsseldorf.

Due to unavailability of part of the data of the JUMAX sample, the final cohort comprised 33 subjects (14 females, mean age 27.5 +/-3 years). Accordingly, to match the number of available subjects from the JUMAX dataset, only the first 33 subjects were used from the IMAX sample (17 females, 27 +/-2.7). For all subsequent analyses, only the first two sessions of both samples were used.

### 2.2 Data acquisition

For both datasets, the data was acquired on a Siemens Magnetom PrismaFit 3-Tesla with a 20-Channel head coil. Structural images were collected using an MP-RAGE sequence (TR=2300ms, TE =2,28ms, TI=900ms, flip-angle=8°) and 1mm voxel size. All RS and NV runs used the same echo planar imaging sequence (TR=719ms, TE=30ms, flip-angle=52°, slices=44, FOV=225×225 mm2) resulting in 2.96×2.96×3 mm voxel size. Data from collaborators at the National University of Singapore were retrieved and structured in the form of a DataLad dataset, a research data management solution providing data versioning, data transport, and provenance capture (Halchenko et al., 2021). Each of the three testing sessions per participant, which were conducted within a seven day period, comprised three NV runs and two RS scans. The order of scans was identical on all three days, starting with a structural scan, followed by 5 functional scans in the order of RS 1, Inscapes, Circus, Indiana Jones and RS 2, with each functional scan lasting for 10 minutes. All movies had been cut to the same length. For RS scans, participants were asked to lay as still as possible and think of nothing in particular, while keeping their eyes open. Instructions for the NV scans were to watch the movies while staying as still as possible. For all scans, participants were asked to not fall asleep during the measurement. Foam wedges were fitted around each subject’s head for comfort and to decrease movement. For all subsequent analyses, only the first two scan sessions and the first RS scan (RS1) of each session were used. The movie clips were presented via a mirror that was mounted on the head coil and the sound was played through headphones.

### 2.3 Stimulus material

Three different movie stimuli with different levels of social content (Inscapes < The Circus < Indiana Jones) were used. Inscapes is a nonverbal, non-social series of animated abstract shapes created by Vanderwal et al. which was looped to match the 10 minutes duration (Vanderwal et al., 2015). The Circus (United Artists Digital Studios, 1928, directed by Charlie Chaplin) is a silent black-and-white. Participants were shown the first 10 minutes of the film which depicts the protagonist being chased by the police and unintentionally causing comic situations during his escape. Indiana Jones and the Temple of Doom (Paramount Pictures, 1984, directed by Steven Spielberg) shows the first 10 minutes of the movie during which the protagonist has to fight off several hitmen who are trying to kill him and finally escapes by taking a plane. The end of the clips used from The Circus and Indiana Jones both coincide with a change of scene in the respective movie itself.

### 2.4 Data preprocessing

Preprocessing of MRI data was performed using fMRIPrep, version 22.0.0 (Esteban et al., 2019). In brief, the T1-weighted volumes were corrected for intensity non-uniformity and skull-stripped. The extracted brain images were then transformed into Montreal Neurological Institute (MNI) space and motion corrected using Advanced Normalization Tools (Avants et al., 2009). The functional data was motion-corrected with MCflirt (Jenkinson et al., 2002) and subsequently co-registered to the native T1-weighted image using boundary based registration with six degrees of freedom from Freesurfer (Greve and Fischl, 2009). Subsequently, an isotropic Gaussian kernel of 6mm FWHM (full-width half-maximum) was applied for spatial smoothing. The images were further regressed out of nuisance signals and bandpass filtered (0.01– 0.1 Hz). Nuisance signals were the global signals extracted within the CSF, the WM, and the whole-brain masks which were regressed from the preprocessed fMRI data for each subject. In addition, the standard six motion parameters and their first temporal derivatives were regressed out.

Subsequently, network functional connectivity (NFC) matrices were constructed for 14 meta-analytical networks, comprising nine to 23 nodes (a detailed description of the networks can be found in the supplements). In short, isotropic 5 mm spheres were created around the local maxima of each meta-analytical network node and only gray matter voxels were included. Using the Junifer toolbox (Synchon Mandal et al., 2023), we extracted the mean time series of each node and computed the Pearson’s correlation coefficient between all node pairs to produce a node times node connectivity matrix for each subject and each condition. The networks cover affective (Amft et al., 2015; Buhle et al., 2014; Liu et al., 2011; Sabatinelli et al., 2011), social (Amft et al., 2015; Bzdok et al., 2012; Caspers et al., 2010), executive(Camilleri et al., 2018; Cieslik et al., 2015; Langner and Eickhoff, 2013; Rottschy et al., 2012), memory (Binder et al., 2009; Spreng et al., 2009) and motor (Witt et al., 2008) functions.

### 2.5 Graph theoretical analyses

Subsequently, graph metrics were derived from the NFC matrices. The fully connected node x node matrices were thresholded at 0.1 to determine the presence or absence of connections (edges) between nodes. Connections above the threshold retained their correlation coefficient, whereas subthreshold edges were assigned values of 0. This thresholding procedure was performed on both positive and negative connections. Five different Graph metrics were extracted from the thresholded NFC matrices using the *NetworkX* toolbox (A Hagberg et al., 2008), including degree centrality, clustering coefficient, betweenness centrality, global efficiency and mean shortest path length. Degree centrality measures the connectedness of each node, computed as the weighted sum of all edges connected to that node. The clustering coefficient for a given node is a measure of local connectedness, measuring the proportion of existing connections out of all possible connections between the nearest neighbors of that node. Betweenness centrality measures the centrality of a node in the network, calculated as the ratio of shortest paths (that is the smallest number of links that need to be traversed to go from one node to another) in the whole graph that pass through that node. The efficiency of a pair of nodes in a graph is the reciprocal of the shortest path distance between these two nodes. The global efficiency of a graph is the average efficiency of all pairs of nodes. Shortest path length denotes the minimum number of nodes that need to be passed through to connect one node to another. Mean shortest path length is the average shortest path length between all nodes of the graph.

### 2.6 Test-retest reliability

The reliability of each graph metric was quantified by calculating the intraclass correlation coefficient (ICC) across these measures derived from the two scans (McGraw and Wong, 1996; Shrout and Fleiss, 1979). A one-way ANOVA was applied to the measures of the two scan sessions across subjects, to calculate between-subject mean square (MSp) and mean square error (MSe). ICC values were then calculated as:

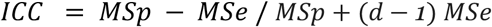

where d is equal to the number of observations per subject. For each graph measure, we calculated reliability at the scan-wise level. Scan-wise reliability estimates the reliability of one score derived from the entire scan session, opposed to calculating one ICC value for the graph metric of each node (Guo et al., 2012; Wang et al., 2017). Here, a single ICC value was calculated for the mean graph metric averaged across all nodes of the network. The reliability results are considered excellent (ICC > 0.8), good (ICC 0.6-0.79), moderate (ICC 0.4-0.59), fair (ICC 0.2-0.39), and poor (ICC < 0.2) (Guo et al., 2012). As negative ICCs are difficult to interpret and reasons for negative values are unclear (Müller and Büttner, 1994), in the following we set negative ICCs to zero (that is completely non-reliable) as has been suggested in previous studies (Braun et al., 2012; Kong et al., 2007; Zhang et al., 2011).

## 3. Results

### 3.1 Reliability of graph metrics in the IMAX sample

We investigated the reliability of five graph measures derived from 14 different networks. For the IMAX sample, we found low to good reliability across networks. Degree centrality, cluster coefficient and efficiency showed a trend towards higher reliability than between centrality and shortest path length.

Degree centrality showed the highest ICC during RS in five (AM, CogAC, MNS, Rew and VigAtt), during Inscapes in three (SM, ER, extDMN), during Circus in four (EmoSF, Empathy, ToM, WM) and during Jones in three (eSAD, Motor, Empathy) networks.

Cluster coefficient showed the highest ICC during RS in four (Rew, Empathy, VigAtt, EmoSF), during Inscapes in three (MNS,ER,extDMN), during Circus in three (AM, SM, ToM) and during Jones in five (CogAC, Motor, EmoSF, eSAD, WM) networks.

Efficiency showed the highest ICC during RS in eight (AM, MNS, CogAC, EmoSF, Rew, eSAD, extDMN, WM), during Inscapes in three (Motor, SM, ER, extDMN), during Circus in three (Empathy, ToM, WM) and during Jones in two (VigAtt, WM) networks.

Between centrality showed the highest ICC during RS in two (EmoSF, Empathy), during Inscapes in five networks (AM, MNS, SM, eSAD, extDMN), during Circus in four (Motor, ER, ToM, WM) and during Jones in three (CogAC, Rew, VigAtt) networks.

Shortest path length showed the highest ICC during RS in four (MNS, EmoSF, eSAD, WM), during Inscapes in four (AM, Motor, Empathy, extDMN), during Circus in two (ER, ToM) and during Jones in four (CogAC, Rew, SM, VigAtt) networks.

### 3.2 Reliability of graph metrics in the JUMAX sample

For JUMAX we found low to excellent reliability across networks. Degree centrality, cluster coefficient and efficiency showed a trend towards higher reliability than between centrality and shortest path length.

Degree centrality showed the highest ICC during RS in nine (AM, CogAC, EmoSF, Empathy, ER, MNS, Motor, VigAtt, WM), during Inscapes in one (eSAD) and during Circus in three (Rew, SM, ToM) networks.

Cluster coefficient showed the highest ICC during RS in five (CogAC, Motor, EmoSF, SM, WM), during Inscapes in two (AM, eSAD), during Circus in four (Rew, ER, ToM, VigAtt) and during Jones in three (MNS, Empathy, extDMN) networks.

Efficiency showed the highest ICC during RS in ten (AM, MNS, CogAC, Motor, EmoSF, Empathy, ER, eSAD, VigAtt, extDMN), during Circus in three (Rew, SM, WM) and during Jones in one (ToM) networks.

Between centrality showed the highest ICC during RS in four (AM, Motor, Rew, ER), during Inscapes in two (Empathy, ToM), during Circus in one (SM) and during Jones in seven (CogAC, EmoSF, MNS, eSAD, VigAtt, extDMN, WM) networks.

Shortest path length showed the highest ICC during RS in nine (AM, MNS, CogAC, EmoSF, Rew, ER, SM, ToM, VigAtt, extDMN), during Circus in two (Empathy, SM) and during Jones in three (Motor, eSAD, WM) networks.

### 3.3 Comparison of the two samples

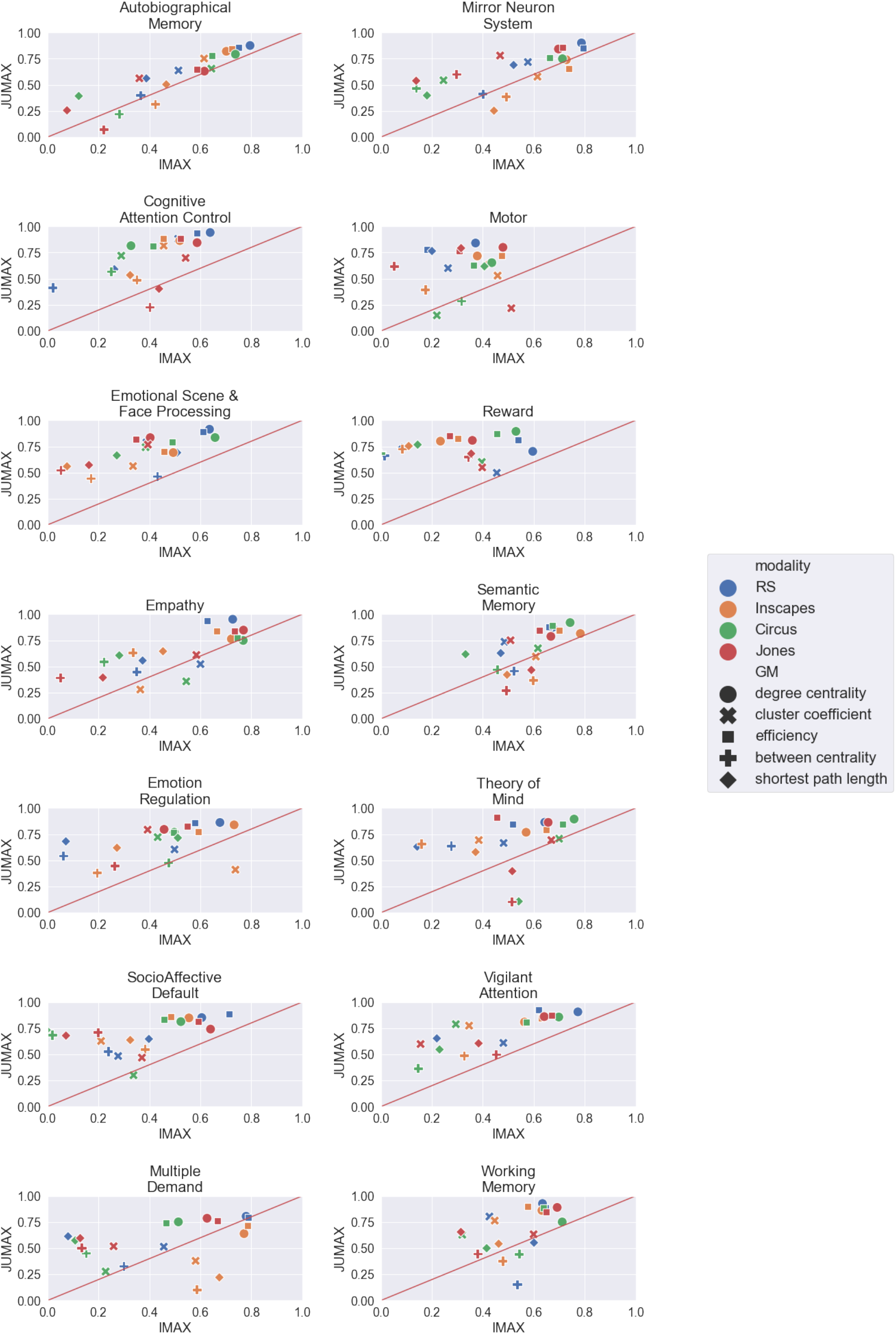

Comparing the results across the two samples, it was evident that the ICC was generally higher in the JUMAX sample than in the IMAX sample. However, in both samples, degree centrality and efficiency tended to show the highest ICCs, followed by cluster coefficient and then by between centrality and shortest path length. The AM, MNS, Empathy and SM networks showed similar results in both samples, while the rest of the networks showed more distinct results. Overall there was not one stimulus which led to more consistent results than other stimuli across the two samples.

## 4 Discussion

The primary goal of this study was to investigate the reliability of NV and RS, across various functional networks. Graph metrics indicate that NV is - in certain conditions - more reliable than RS, consistent with previous results from Wang et al. 2017 (Wang et al., 2017). However, our results demonstrated that this effect is dependent on a variety of factors. Firstly, the choice of the NV stimulus impacts the reliability of a given graph metric. Secondly, the effect of NV stimuli varies across cohorts. Thirdly, the increase in reliability is not uniform across the brain, but varies between different functional networks.

### NV vs RS

Starting from observations indicating that graph metrics extracted from RS fMRI can be used to investigate abnormalities in brain organization (Petrella, 2011; Wu et al., 2009), researchers have focused on investigating the reliability with which these graph metrics can be extracted. With ongoing efforts to use characteristic abnormalities to successfully detect and track neurological diseases, it will be crucial to increase reliability as much as possible. Therefore, researchers have shifted to extracting graph metrics from other modalities than RS such as task-based fMRI (Aron et al., 2006; Cao et al., 2014) or NV (Rikandi et al., 2022; Zhang and Liu, 2021). In contrast to task-free RS, these modalities place a constraint on the participant which might reduce variability that is otherwise induced by spontaneous thoughts (Finn et al., 2017; Hasson et al., 2010; Vanderwal et al., 2017). Our results confirmed the notion that behavioral constraints can prove to be beneficial to increase reliability over unconstrained RS. In multiple networks, NV stimuli increased reliability of one or more graph metrics in comparison with RS (Fig. 1, Fig. 2). Furthermore, this improvement of reliability is observable across networks dealing with affective, social, executive, memory and motor functions, indicating that NV increases engagement not only in sensory, but also in higher order networks. On the other hand, our results also showed that in many instances RS was more reliable than NV, which is in line with previous studies that showed that NV does not unconditionally increase reliability (Hlinka et al., 2022; Zhang et al., 2022). Nevertheless, these results, in our opinion, encourage the use of NV to improve reliability as NV increased reliability over RS drastically in certain cases. But rather than viewing NV as a one-fits-all tool, our findings further underline the importance of using specific NV stimuli (and brain networks) for a specific purpose.

**Figure 1.**
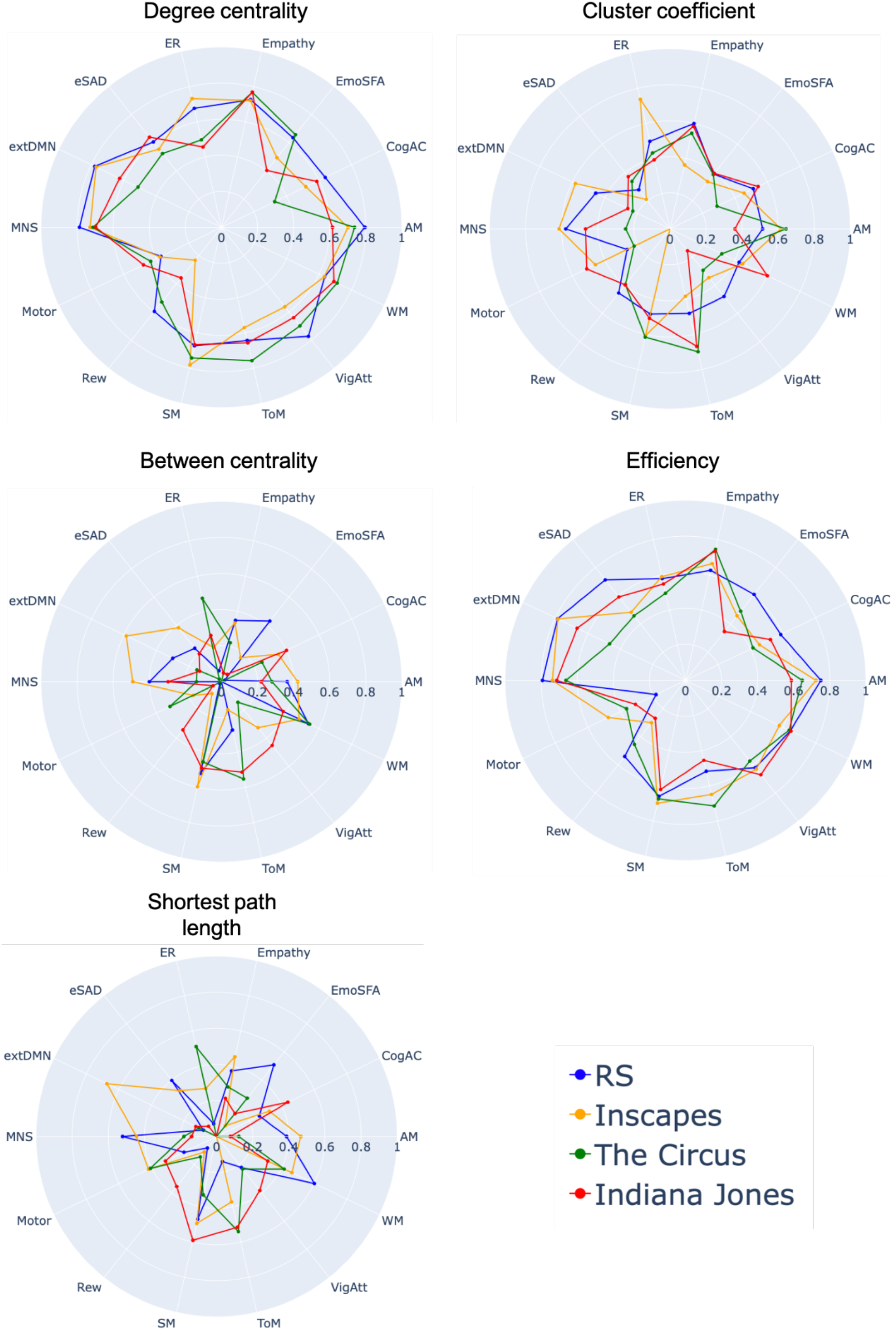
ICC of graph metrics across the 14 networks in the IMAX sample. Graph metrics are shown for the RS scan and three different movies. ICC values below zero are not depicted. (AM =Autobiographical Memory, CogAC = Cognitive Attention Control,eMDN=extended Multiple Demand Network, EmoSF= Emotional Scene and Face Processing, ER = Emotion Regulation, eSAD=Extended Social-affective Default, MNS = Mirror Neuron System, Rew = Reward, SM = Semantic Memory, ToM = Theory of Mind, VigAtt= Vigilant Attention, WM = Working memory)

**Figure 2.**
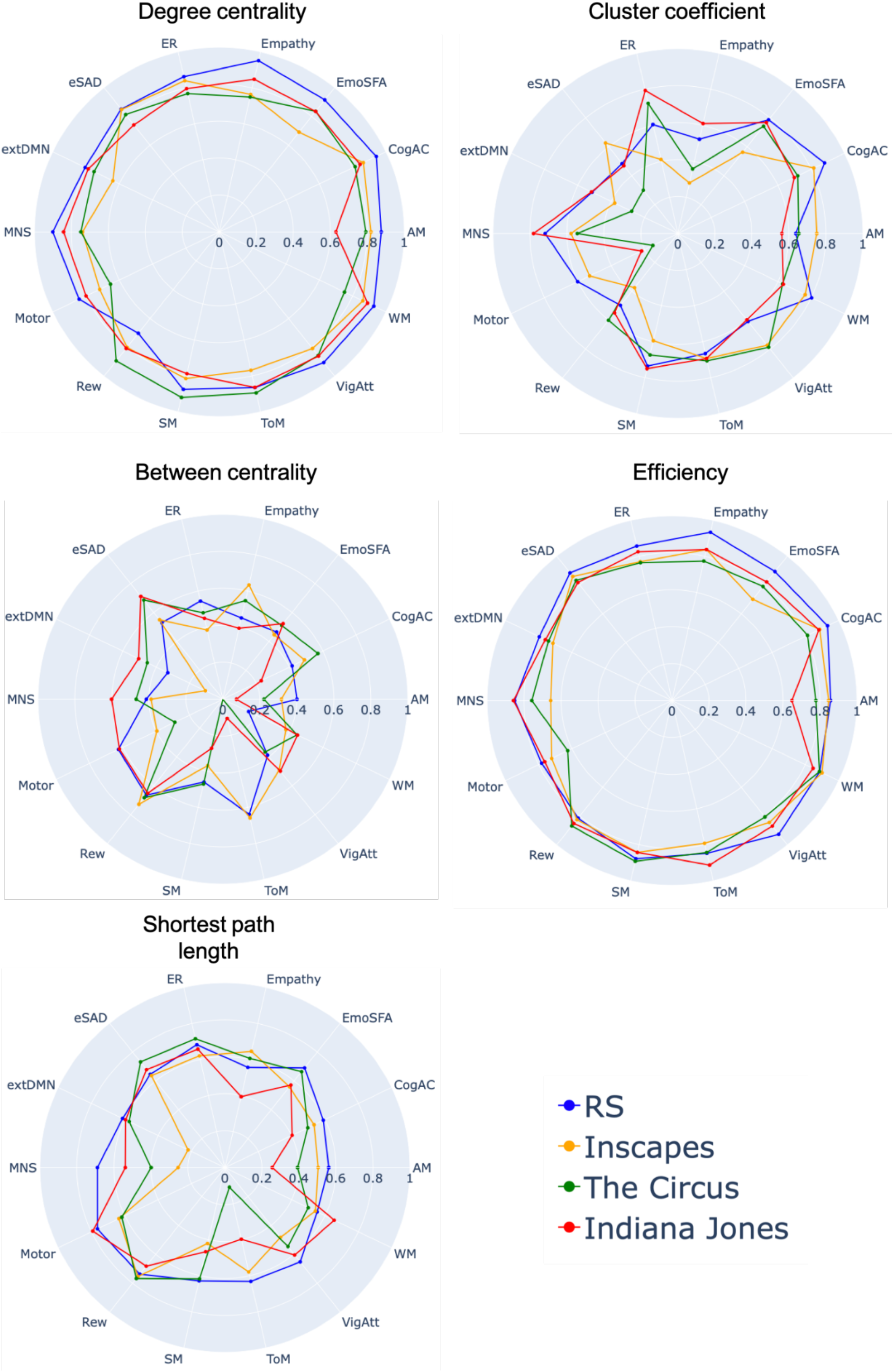
ICC of graph metrics across the 14 networks in the JUMAX sample. Graph metrics are shown for the RS scan and three different movies. ICC values below zero are not depicted. (AM =Autobiographical Memory, CogAC = Cognitive Attention Control,eMDN=extended Multiple Demand Network, EmoSF= Emotional Scene and Face Processing, ER = Emotion Regulation, eSAD=Extended Social-affective Default, MNS = Mirror Neuron System, Rew = Reward, SM = Semantic Memory, ToM = Theory of Mind, VigAtt= Vigilant Attention, WM = Working memory)

The observed reliability in our study matches results from previous studies investigating graph metrics extracted from RS and NV (Braun et al., 2012; Cao et al., 2014; Wang et al., 2017). However, in contrast to Wang et al (Wang et al., 2017) we showed that NV does not generally improve reliability of graph metrics, but that its effect varies across networks, stimuli and graph metric.

### Variance across cohorts

One of the advantages of using NV stimuli is that they are easier to share across multiple sites than traditional tasks (DuPre et al., 2020; Eickhoff et al., 2020). By combining data from a multitude of studies using the same NV stimuli, one can not only achieve large sample sizes, but also place the same behavioral constraint on all subjects across sites. However, several studies have suggested that cultural differences between movies (and/or cohorts) might hinder generalizability (DuPre et al., 2020; Eickhoff et al., 2020; Hasson et al., 2010). In this study, we compared an asian and a european cohort that were subjected to the same three NV stimuli. In both samples, NV stimuli increased reliability of graph metrics in comparison with RS. However, we did not observe that the same combination of stimulus, network and graph metric led to improved reliability over RS in both samples (Fig. 3). Although some of the networks (AM, MNS, Empathy and SM) show similar trends, it is not generally the case that results from both samples are highly overlapping. These differences might have been driven by the different cultural backgrounds of the participants. The appreciation of a film is culturally specific (Saarimäki, 2021) and likely different between the european and asian cohorts. Several studies have demonstrated cultural differences in the perception of faces (Adams et al., 2010; Goh et al., 2010; Harada et al., 2020), a factor that is especially relevant for the NV stimuli Circus and Jones during which a variety of different faces are depicted. Related, in a study from Sneddon et al., 2011 (Sneddon et al., 2011) participants from Northern Ireland, Serbia, Guatemala and Peru showed systematic differences in their rating of positive and negative emotions being displayed in twelve short movie clips. Our study provides further evidence for the notion that future studies should take into account cultural differences between cohorts when selecting a movie stimulus.

### Variance across networks and stimuli

In our analysis, we employed meta-analytically defined networks that represent the most likely core nodes of a given brain function. Alternatively to approaches where the effect of NV is considered from a whole brain perspective, we here investigated how NV engages different networks. Similar to previous studies, we observed that the effect of different NV stimuli varies across different networks (Finn and Bandettini, 2020; Kröll et al., 2023; Wang et al., 2017) and reliability of graph metrics was not unconditionally increased over RS. One of the advantages of NV is the possibility to more effectively engage brain networks of interest, in comparison with RS (Eickhoff et al., 2020; Guo et al., 2015). Intuitively, one would expect that a network responsible for the processing of emotions is differently engaged by an emotional clip than e.g. the motor network. This effect can also be seen in our results as different networks exhibit varying reliabilities in response to the same stimulus To analyze the effect of the chosen movie stimulus on the reliability of a given graph metric, we employed three movies with different levels of social content. Various studies have shown that different NV stimuli can lead to significantly different results. Finn et al., 2021 reported that FC derived from different movies varied in its ability to accurately predict emotion and cognition scores (Finn and Bandettini, 2020). Similarly, Gal et al., 2022 showed that the accuracy with which task-activation maps could be predicted differed between FC derived from Hollywood NV stimuli and independent NV stimuli (Gal et al., 2022). Our results extend these findings by showing that NV stimuli also divert in their impact on the reliability of extracted graph measures. Previous studies have shown that reliability is strongly dependent on attention (Ki et al., 2016) and several studies have suggested that NV stimuli with social content are best suited to engage participants and keep their attention over a longer period of time (Finn and Bandettini, 2020; Saarimäki, 2021; Schaefer et al., 2010). In line with that, we observed a tendency that the two more social stimuli, Circus and Jones, more frequently led to improvement than the abstract movie Inscapes. However, in the majority of cases, reliability was higher for graph metrics extracted from RS than these extracted from NV. This was somewhat unexpected since RS is generally seen as an unconstrained state and one would therefore expect more variability between sessions than for more constrained states like NV. Several factors might have led to the relative decrease in reliability for NV. Firstly, familiarity with a given movie might have played a role as multiple studies have shown that expected stimuli reduce the neuronal response (Alink et al., 2010; Koster-Hale and Saxe, 2013). The sessions for both datasets were conducted within a week and therefore participants will be familiar with the movie during the second session. This effect might have induced variability for the NV conditions, while RS on the other hand has been shown to remain stable across sessions (Mason et al., 2007; Wang et al., 2011). Secondly, some of the networks employed here (AM, SM and eSAD) are overlapping with the default mode network which is linked to intrinsically oriented functions, rather than the processing of external stimuli (Golland et al., 2007; Hasson et al., 2004). This may plausibly lead to decreased reliability of NV in comparison with RS, in these networks. These results emphasize that future studies should carefully consider which combination of graph metric, stimulus and network is suited for the research question at hand. Using purpose-built movies, such as emotionally salient clips for patients with depression (Guo et al., 2016), in combination with the functionally involved network will help improve reliability and advance the characterization of disease specific alterations in the brain.

## 4 Limitations

While the current study sheds new light onto the reliability of NV in comparison with RS, it comes with some limitations. Firstly, the reliability of graph metrics is strongly influenced by the choice of the applied preprocessing (Andellini et al., 2015). In this study, we applied motion correction and regressed out WM and CSF signals, as has been done in most previous studies (Braun et al., 2012; Cao et al., 2014; Wang et al., 2017). On top of that, we here applied basic (that is only removing the mean signal of the whole brain) global signal regression. There is an ongoing debate of whether or not to apply global signal regression, with some studies claiming that it introduces spurious anti-correlations while other reports suggest that these anti-correlations are true negative connections (Liang et al., 2012; Murphy et al., 2009). However, a review by Andellini et al., 2015 found no significant differences between the reliability of data with and without the inclusion of global signal regression across five studies (Andellini et al., 2015). Secondly, we here considered both, negative and positive connections, with the assumption that both are true representations of connectivity. However, several papers have indicated that negative correlations should be evaluated with care since they tend to reduce test-retest reliability (Andellini et al., 2015; Schwarz and McGonigle, 2011; Wang et al., 2011). Therefore, the reliability of single graph metrics in our study might have been decreased by the inclusion of negative connections. Thirdly, our results are based on weighted adjacency matrices, because they better characterize the underlying connectivity by considering connectivity strength. However, previous studies have suggested that binarized adjacency matrices may lead to higher reliability (Andellini et al., 2015; Wang et al., 2011). Nevertheless, we think that using weighted adjacency matrices is preferable, especially for clinical studies where subtle changes in connectivity might help to identify disease specific alterations.

## Conclusion

NV has been suggested to improve the reliability of graph based measures in comparison with RS. Our findings extend the current knowledge by investigating this effect in different networks, with multiple NV stimuli and in two different cohorts. We demonstrate that the potential increase in reliability is dependent on the chosen NV stimuli and varies between functional networks. Furthermore we suggest that cultural differences should be considered when sharing NV stimuli across sites. Our study supports the use of NV to increase reliability of graph metrics, but emphasizes the need to carefully select the appropriate stimulus and network for the research question at hand.

## Acknowledgment

This work was supported by the European Union’s Horizon 2020 Research and Innovation Programme under grant agreement no. 945539 (HBP SGA3), and the Deutsche Forschungsgemeinschaft (491111487). We also acknowledge the funding support from Yong Loo Lin School of Medicine, National University of Singapore (J.H.Z), the Duke-NUS Medical School Signature Research Program Core Funding (J.H.Z.), and Ministry of Education, Singapore (MOE-T2EP40120-0007, J.H.Z), and Far East Organization (E-546-00-0398-01, MWLC.).

